# The cortical N1 response to balance perturbation is associated with balance and cognitive function in different ways between older adults with and without Parkinson’s disease

**DOI:** 10.1101/2022.02.08.479608

**Authors:** Aiden M. Payne, J. Lucas McKay, Lena H. Ting

## Abstract

Mechanisms underlying associations between balance and cognitive impairments in older adults with and without Parkinson’s disease are poorly understood. Balance disturbances evoke a cortical N1 response that is associated with both balance and cognitive abilities in unimpaired populations. We hypothesized that the N1 response reflects neural mechanisms that are shared between balance and cognitive function, and would therefore be associated with both balance and cognitive impairments in Parkinson’s disease. Although N1 responses did not differ at the group level, they showed different associations with balance and cognitive function in the Parkinson’s disease vs. control groups. In the control group, higher N1 amplitudes were correlated with lower cognitive set shifting ability and lower balance confidence. However, in Parkinson’s disease, narrower N1 widths (i.e., shorter durations) were associated with greater parkinsonian motor symptom severity, lower balance ability and confidence, lower mobility, and lower overall cognitive function. Despite different relationships across populations, the present results suggest the N1 response reflects neural processes related to both balance and cognitive function. A better understanding of neural mechanisms linking balance and cognitive function could provide insight into associations between balance and cognitive decline in aging populations.

## INTRODUCTION

### Assessing cortical activation during balance recovery behavior may provide insight into relationships between balance and cognitive impairments with aging and Parkinson’s disease

A systematic review and meta-analysis found that lower scores on global measures of cognitive function, and executive function in particular, predict future falls in otherwise healthy older adults (Muir et al. 2012). Global measures of cognitive function (Kim et al. 2013) and executive function (Hausdorff et al. 2006; Mak et al. 2014) likewise predict future falls in Parkinson’s disease. The mechanisms linking balance and cognitive impairments are unclear but may be reflected in cortical activation during balance recovery. Although a balance disturbance initially evokes an automatic brainstem-mediated balance-correcting motor reaction, cortical contributions to balance recovery can follow as needed (Jacobs and Horak 2007a). Older adults, and particularly people with Parkinson’s disease, generally show increased cortical activity compared to young adults for similar performance levels of walking and balance tasks (Stuart et al. 2018), which may reflect cognitive engagement to compensate for impaired automatic control of balance (Petzinger et al. 2013; Wu et al. 2015a), providing a potential opportunity for cognitive impairment to influence balance control. Increased recruitment of cortical activity for motor tasks with aging can be observed at relatively younger ages for more complex motor tasks (Nobrega-Sousa et al. 2020), and individuals with greater cortical recruitment in simple single task conditions have greater difficulty performing more complex tasks that push the limits of compensatory recruitment of their remaining executive control resources (Hawkins et al. 2018; Palmer et al. 2021). Cortical responses evoked in cognitive tasks have been linked to cognitive impairment in a variety of studies (reviewed in Seer et al. 2016 and Wang et al. 2020), but in the present study we investigate a cortical response evoked during balance recovery that has been associated with both balance and cognitive abilities in healthy populations, and may therefore provide insight into relationships between balance and cognitive impairments in Parkinson’s disease.

### A balance disturbance evokes a fast cortical response that has been associated with both balance ability and cognitive processing in healthy populations

A sudden balance disturbance evokes a cortical “N1” response peak at 100-200 ms in electroencephalography (EEG) activity that has been localized to the supplementary motor area in healthy young adults using a single-source assumption (Marlin et al. 2014; Mierau et al. 2015). However, time-frequency analyses suggest the anterior cingulate cortex, sensorimotor areas, and parietal cortex synchronize with the supplementary motor area during the N1 response (Peterson and Ferris 2018; 2019; Varghese et al. 2019), suggesting a network of cortical areas may contribute to the N1 response. The supplementary motor area is thought to mediate interactions between motor and cognitive processes through its connections to these other cortical areas (Goldberg 1985), which could be reflected in the N1 response. In young adults the cortical N1 is larger in individuals with lower balance ability (Payne and Ting 2020a) and on trials that include compensatory stepping behaviors (Payne and Ting 2020c; Solis-Escalante et al. 2020), possibly reflecting compensatory cortical engagement in balance recovery. The cortical N1 is also influenced by cognitive processing in young adults, becoming smaller when attention is directed away from balance recovery by a dual task paradigm (Little and Woollacott 2015; Quant et al. 2004b), and larger when perturbations are perceived to be more threatening (Adkin et al. 2008; Mochizuki et al. 2010) or less predictable (Adkin et al. 2008; Mochizuki et al. 2010). While these within-subjects studies demonstrate a causal influence of changes in the availability or allocation of cognitive processing resources on the N1 response, our prior work was the first to assess whether individual differences in cognitive ability were reflected in the N1 response. Specifically, we found that otherwise healthy older adults who were slower in the executive function of cognitive set shifting also displayed larger cortical N1 amplitudes, stiffer balance recovery behavior, and increased antagonist muscle activity (Payne et al. 2021), further implicating the neural processes underlying the N1 in the relationship between balance and cognitive problems with aging. While studies in older populations have been limited, older adults generally have smaller and later perturbation-evoked N1s (Duckrow et al. 1999; Ozdemir et al. 2018), with changes in temporal characteristics including the appearance of multiple component peaks in some individuals with reduced mobility (Duckrow et al. 1999). We now investigate group- and individual-differences the cortical N1 responses in populations of older adults with and without Parkinson’s disease.

### Parkinson’s disease affects several factors known to influence the cortical N1, but it is unknown whether the N1 is altered in Parkinson’s disease

The N1 depends on attention to balance control (Little and Woollacott 2015; Quant et al. 2004b), which is increased Parkinson’s disease (Petzinger et al. 2013; Wu et al. 2015a). N1 amplitude also depends on the perceived threat of a balance disturbance (Adkin et al. 2008; Mochizuki et al. 2010), and fear of falling is common in Parkinson’s disease (Grimbergen et al. 2013). Additionally, N1 amplitude in younger adults is associated with lower balance ability (Payne and Ting 2020a), a hallmark of Parkinson’s disease (Bloem 1992; Grimbergen et al. 2004; Koller et al. 1989). Further, in older adults, N1 amplitude is associated with lower cognitive set shifting ability and greater antagonist muscle activity (Payne et al. 2021), both of which are associated with balance impairment in Parkinson’s disease (Lang et al. 2019; McKay et al. 2018). All of these associations in unimpaired populations suggest the N1 would be larger in Parkinson’s disease, related to greater cortical engagement to compensate for balance impairments, but there are also reasons to suspect the N1 might be reduced in Parkinson’s disease. The supplementary motor area is implicated as a major contributor to the cortical N1 response (Marlin et al. 2014; Mierau et al. 2015) and is the cortical node of the basal ganglia thalamocortical “motor circuit” that is impaired in Parkinson’s disease (Albin et al. 1989; Alexander and Crutcher 1990; Alexander et al. 1991; Alexander et al. 1986). Further, the N1 resembles the more widely studied error-related negativity (Payne et al. 2019b), which is reduced in amplitude in Parkinson’s disease (Seer et al. 2016). The error-related negativity is evoked by mistakes in cognitive tasks, depends on dopamine (de Bruijn et al. 2004; de Bruijn et al. 2006; Zirnheld et al. 2004) and connections to the basal ganglia (Ullsperger et al. 2014). If the N1 shares underlying mechanisms with the error-related negativity, we would expect a reduced amplitude in Parkinson’s disease. A brief report on balance N1s in mild Parkinson’s disease showed multiple component peaks (Dimitrov and Gatev 2001) resembling N1s in older adults without Parkinson’s disease (Duckrow et al. 1999), but did not include a control group or measures of balance or cognitive function. Here we compare cortical N1s between people with and without Parkinson’s disease, and test for associations with various measures related to balance and cognitive function.

### We hypothesized that the N1 response reflects neural processing related to both balance and cognitive function, and would therefore be altered in Parkinson’s disease in association with balance and cognitive impairments

We evoked the cortical N1 response using unpredictable forward and backward translations of the support surface. We assessed the amplitude and temporal characteristics of the cortical N1, including the amplitude, latency, and width of the evoked peak. We used multiple measures of balance and mobility, including the clinical miniBESTest (Leddy et al. 2011), the Timed Up and Go test (Beauchet et al. 2011), and measures of cognitive function, including the Montreal Cognitive Assessment (Nasreddine et al. 2005) and the Trail Making Test (McKay et al. 2018; Sanchez-Cubillo et al. 2009). Although we did not find group-level differences in the cortical N1 response amplitude, latency, or width, associations between the cortical N1 and the various balance- and cognitive-related metrics differed between the groups with versus without Parkinson’s disease.

## MATERIALS AND METHODS

### Study populations

#### Participants

Sixteen older adults with Parkinson’s disease (PD, N=16, age 69±7, 4 female) and nineteen older adults without Parkinson’s disease (noPD, N=19, age 71±6, 6 female) are included in analyses after exclusion of four participants as detailed below. Written consent was obtained from all participants after a detailed explanation of the protocol according to procedures approved by the Emory University Institutional Review Board. Different analyses have been previously reported in the noPD control group (Payne et al. 2021).

#### OFF-medications

Individuals with PD participated in the experiment OFF their dopamine medications, practically defined as a minimum of 12 hours after their last dose of dopaminergic medication for PD. Each participant’s neurologist was consulted and signed an OFF-medication clearance form before they were asked to withhold their medications for the purpose of this study. All clinical and behavioral measures were collected during the same OFF-medication session, with disease duration and compatibility with inclusion/exclusion criteria additionally verified in patient clinical records when available.

#### Participant recruitment

Participants were recruited from the community surrounding Emory University and the Emory Movement Disorders clinic through flyers, outreach events, word of mouth, and databases of prior participants from collaborating groups. Adults over age 55 were screened for the following inclusion criteria: vision can be corrected to at least 20/40 with glasses, no history of stroke or other neurological condition (except for PD), no musculoskeletal conditions or procedures that cause pain or limit mobility of the legs, ability to stand unassisted for at least 15 minutes, and cognitive ability to consent. Potential participants were excluded for prior experience on the perturbation platform, present cholinergic medications, or lack of neurologist’s approval to withhold dopaminergic medications. Participants with PD were recruited first, and then the older adult control participants were recruited to maintain similar age and sex distributions between groups.

Four participants with PD were excluded after partial or complete participation in the study, resulting in the reported N=16 after an initial recruitment of N=20. Two were excluded due to either a brain tumor or severe peripheral neuropathy of the legs noted in their clinical record. One was excluded due to failure to save the EEG data. One was unable to tolerate being OFF-medication and opted to leave prior to the balance perturbations.

### Experimental protocol and data collection

#### Parkinson’s disease motor symptom severity

The motor subscale of the International Parkinson and Movement Disorder Society’s Unified Parkinson’s Disease Rating Scale (MDS-UPDRS III) was used to assess the severity of motor impairment in participants with PD (Goetz et al. 2007). The test was administered by AMP, who is certified by the Movement Disorders Society, and filmed for subsequent scoring by a practicing neurologist. Postural instability/gait difficulty subscores were determined from the items of the MDS-UPDRS III (Stebbins et al. 2013) and included in analyses. Hoehn & Yahr (H&Y) stage (Goetz et al. 2004), a 5 point rating scale of PD severity focused on postural instability, was determined from the recorded videos by a neurologist and included in analyses.

#### Parkinson’s disease duration

Participants with PD were asked to report the number of years since PD diagnosis at the time of participating in the study, and this was verified in the clinical record when possible.

#### Balance ability

The mini Balance Evaluation Systems Test (miniBESTest, www.bestest.us) was used as a measure of balance ability (Leddy et al. 2011; Lofgren et al. 2017; Magnani et al. 2020), which assesses anticipatory postural control, reactive postural control, sensory orientation, and dynamic gait. For items that scored the left and right sides separately, only the lower of the two scores was considered for a maximum possible score of 28 (Lofgren et al. 2017).

#### Balance Confidence

The Activities-Specific Balance Confidence (ABC) Scale (Powell and Myers 1995) was used to assess balance confidence. This survey consists of sixteen items describing different situations that might lead to a loss of balance. For each item, participants are asked to indicate their confidence that they would not lose their balance in a particular setting by answering with a percentage between 0-100%. The average score across the 16 items is reported as the total score. This measure predicts falls in healthy older adults (Cleary and Skornyakov 2017) and differs between recurrent and non-recurrent fallers with Parkinson’s disease (Mak et al. 2014), and is included as a potential between-subjects assessment of previously established within-subjects effects of perceived threat on the perturbation-evoked N1 response (Adkin et al. 2008).

#### Mobility

The Timed Up and Go (TUG) test (Beauchet et al. 2011) was administered within the miniBESTest, and additionally scored in more detail than considered within the miniBESTest. Participants begin seated in a chair with arms in their lap, and when told to “Go,” must get up, walk at their comfortable speed across the lab, around a cone, and come back to a seat in the starting chair. This test is timed, and then repeated with a secondary task of counting backward by 3’s out loud. While the miniBESTest only scores this item categorically, based on whether participants were able to complete the dual task condition, and if so, whether it resulted in more or less than a 10% reduction in speed, we included additional continuous measures in our analyses. Specifically, we included the TUG single task time (TUG-ST), dual task time (TUG-DT), and dual task interference (DTI) calculated as the difference between the single and dual task times divided by the single task time and multiplied by 100 (Kelly et al. 2010; Palmer et al. 2021). A more negative value for DTI indicates a greater reduction in speed during the dual task condition. TUG-ST is an indicator of walking mobility, which has previously been associated with differences in the temporal features of the perturbation-evoked N1 response in older adults (Duckrow et al. 1999). DTI can be an indicator of cognitive engagement in balance control, and we have previously demonstrated that DTI during walking is associated with sensorimotor-prefrontal beta coherence during the single-task perturbation condition in a different cohort of healthy older adults (Palmer et al. 2021).

Two individuals with Parkinson’s disease were unable to complete the TUG-ST or TUG-DT due to mobility impairments including freezing of gait, and an additional two individuals were able to complete TUG-ST but not TUG-DT. These individuals are therefore excluded from the corresponding continuous measures but could be appropriately scored on the miniBESTest.

#### Global cognition

The Montreal Cognitive Assessment (MoCA, www.mocatest.org) was used as a global measure of overall cognitive ability, including executive function, attention, and memory (Hoops et al. 2009; Nasreddine et al. 2005). This measure is included based on prior findings linking global measures of cognitive function to falling in older adults with (Kim et al. 2013) and without (Muir et al. 2012) Parkinson’s disease.

#### Cognitive set shifting ability

The set shifting ability score was measured as the difference in time to complete Part B minus Part A of the Trail Making Test (McKay et al. 2018; Payne et al. 2021; Sanchez-Cubillo et al. 2009), where a longer time to complete Part B compared to Part A indicates lower cognitive set shifting ability. This test is frequently used as a measure of executive function that has been associated with falling in people with (McKay et al. 2018) and without (Muir et al. 2012) Parkinson’s disease, and we have previously demonstrated that cognitive set shifting ability is associated with individual differences the cortical N1 response amplitude in the present healthy cohort (Payne et al. 2021).

#### Education

Years of education was self-reported and is included in analyses based on its established relationship to scores on the Montreal Cognitive Assessment (Rossetti et al. 2011) and the Trail Making Test (Giovagnoli et al. 1996; Tombaugh 2004). However, it Is argued that years of education is a poor proxy for intelligence, which better accounts for its association to standard cognitive tests (Steinberg et al. 2005).

#### Perturbations

A series of 48 translational support-surface perturbations of unpredictable timing, direction, and magnitude were delivered during quiet standing (Payne et al. 2021). Perturbations were delivered using a custom-designed perturbation platform (Factory Automation Systems, Atlanta, GA) using 2 brushless AC motors controlled by two servo controllers and a motion controller (BSM80N-375AF and NSB002-501 from ABB Motors and Mechanical Inc., Fort Smith, AR). Perturbations consisted of forward and backward perturbation directions of three magnitudes. The low magnitude (0.15 g, 11.1 cm/s, 5.1 cm) was identical across participants, while the medium (0.21-0.22 g, 15.2-16.1 cm/s, 7.0-7.4 cm) and high (0.26-0.29 g, 19.1-21.0 cm/s, 8.9-9.8 cm) magnitudes were adjusted according to participant height as previously described (Payne et al. 2021) to account for the effect of height on the cortical responses (Payne et al. 2019a) and to ensure that the more challenging perturbations were mechanically similar across different body sizes. Perturbation characteristics for an example participant are shown in Figure 1.

**Figure 1.**
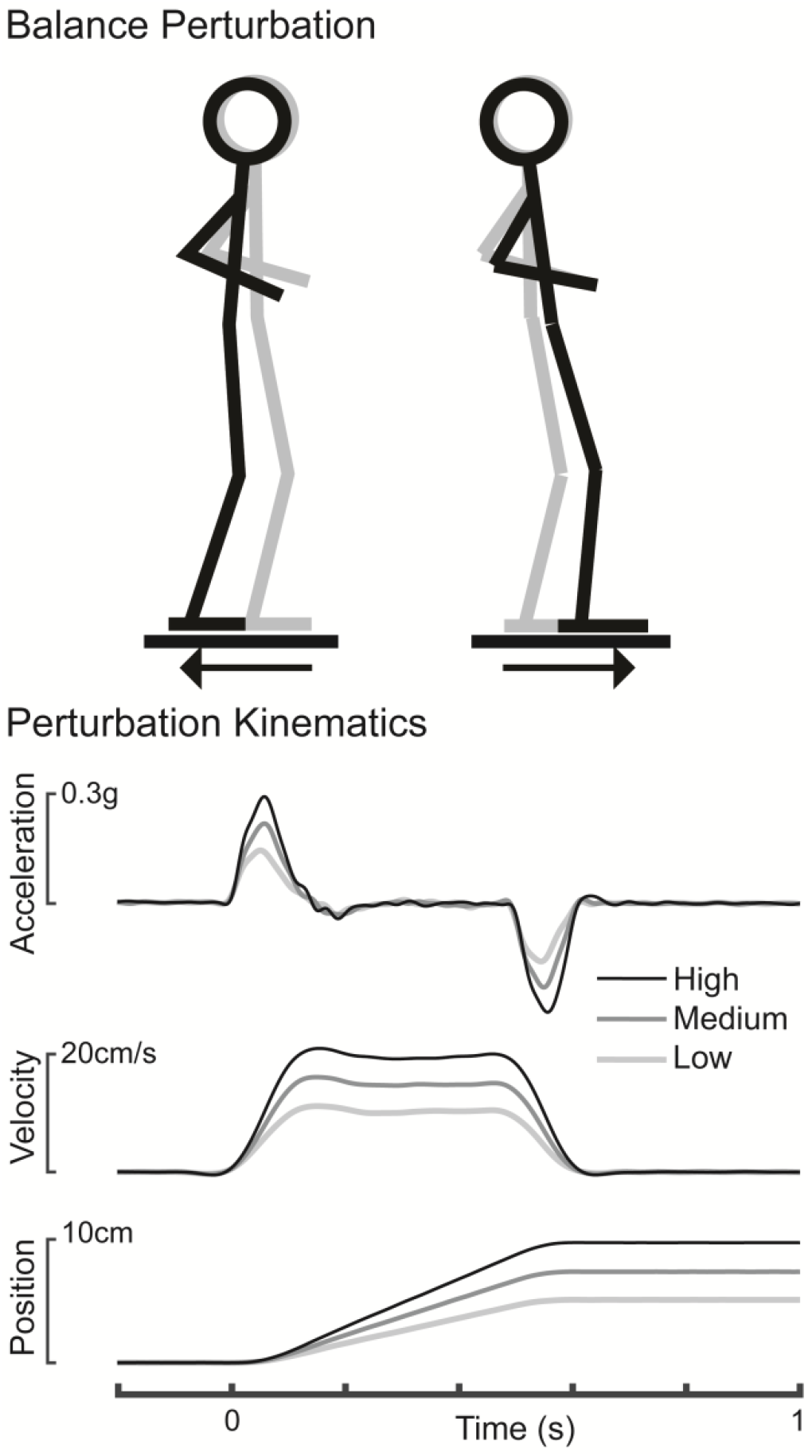
Balance perturbations. A schematic shows the support-surface perturbation along with the measured perturbation kinematics for an example participant (194 cm in height).

To minimize effects of fatigue, a 5-minute break was enforced halfway through the perturbation series, or more frequently without limitations if requested. Excluding rest breaks, the duration of the perturbation series was 21±2 minutes (PD: 20±1 minutes; noPD: 21±2 minutes). Inter-trial-intervals, measured between perturbation onsets, excluding rest breaks longer than a minute, were 23±12 seconds (PD: 23±13 s; noPD: 23±12 s).

Recording artifacts were minimized during data collection by ensuring that perturbations were only initiated during a relatively quiescent baseline in the live electroencephalography (EEG) data based on visual inspection. Participants were instructed maintain their arms crossed across their chest, focus their vision on a poster of a mountain landscape 4.5 m ahead, and to do their best to recover balance without taking a step. Trials in which steps were taken (8% of all trials; PD: 9%; noPD: 8%) were excluded from analysis.

#### Cortical activity

EEG data were collected during the perturbation series as previously described (Payne et al. 2021). Thirty-two active electrodes (ActiCAP, Brain Products, Germany) were placed according to the international 10-20 system, except for two reference electrodes placed on the mastoid bones behind the ears. Electrodes were prepared with conductive gel (SuperVisc 100 gr. HighViscosity Electrolyte-Gel for active electrodes, Brain Products) using a blunt-tipped syringe that was also used to abrade the skin to reduce impedances. Analyses were focused at the Cz electrode, where the N1 response was the largest. Impedances at Cz and mastoid electrodes were generally below 10 kOhm prior to data collection.

Electrooculography (EOG) data were collected to enable subtraction of eye-related artifacts. Bipolar passive electrodes (E220x, Brain Products) were prepared with abrasive gel (ABRALYT HiCl 250 gr., High-chloride-10% abrasive electrolyte gel, Brain Products) and placed above and below the right eye and referenced to a similar electrode on the forehead. EEG and EOG data were sampled at 1000 Hz on an ActiCHamp amplifier (Brain Products) with a 24-bit A/D converter and an online 20 kHz anti-aliasing low-pass filter.

EEG and EOG data were filtered between 1 Hz and 25 Hz using sixth-order zero-lag Butterworth filters. This filtering is consistent with prior time-domain analyses of the N1 response (Marlin et al. 2014; Mierau et al. 2015) in that there is negligible power reduction in the theta frequency (3-8 Hz) range, which contains the N1 response. Cz data were then re-referenced to the mastoids and epoched between 400 ms before to 2000 ms after perturbation onset (defined based on recorded platform acceleration, Figure 1). Blink and vertical eye movement artifacts were subtracted using a serial regression and subtraction approach (Gratton et al. 1983) as previously described (Payne et al. 2019a). Because small translational perturbations applied at the feet do not result in substantial head movement until after the N1 response (Payne et al. 2019a), and because perturbations were only initiated when the participant and EEG baseline had been steady for several seconds, no potential movement artifacts were observed during the time window of the N1 response. Accordingly, no trials were rejected on the basis of visual inspection.

Cz epochs were then averaged across non-stepping trials within each individual and baseline subtracted using a baseline of 50-150 ms before perturbation onset. Cortical N1 response amplitudes and latencies were quantified as amplitude and latency of the most negative point between 100-200 ms after perturbation onset in the subject-averaged EEG waveform at Cz. Because the waveform shape differed to a large extent between individuals, often containing multiple peaks, but not consistently enough to enable measurement of a distinctly identifiable additional peak across individuals (Figure 2 CD), cortical N1 width was assessed using the full-width half-maximum. Specifically, the duration that the N1 response continuously maintained at least half of its most negative amplitude was measured for each individual.

**Figure 2.**
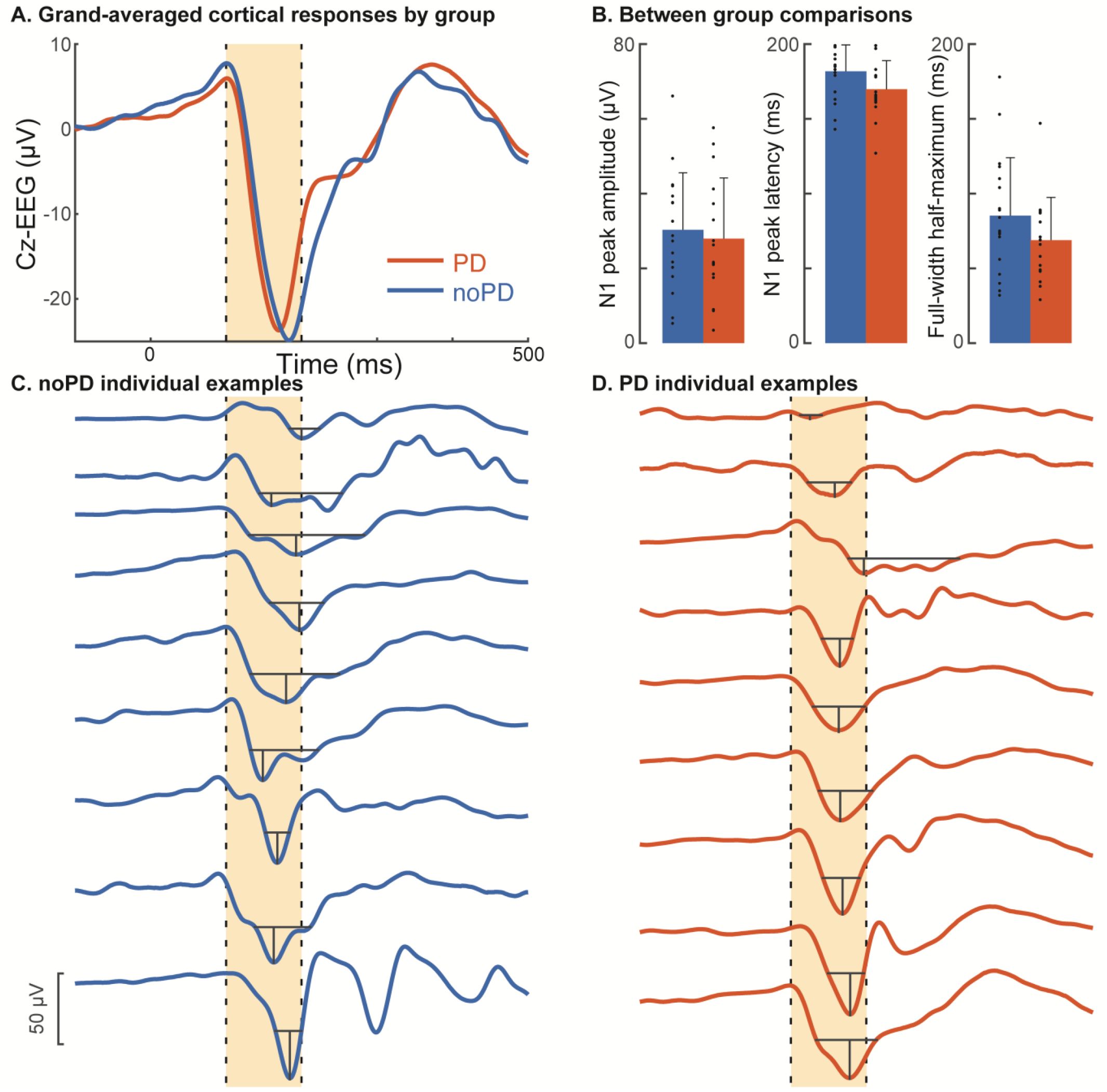
Differences in N1 responses between noPD and PD groups. (A) Grand-averaged cortical responses for each participant group. The yellow shaded region indicates the 100-200 ms window in which N1 peak amplitudes and latencies were quantified. (B) Bar plots show means and standard deviations of N1 peak amplitudes, latencies, and widths by group. Dots show individual data points. The lower panels show individual examples of subject-averaged cortical N1 responses at Cz in (C) the noPD control group and (D) the PD group. N1 peak amplitudes and latencies are indicated by vertical black lines and the duration of the full-width half-maximum is indicated by horizontal black lines.

### Statistical Analyses

#### Between-group comparisons

Two-tailed t-tests were used to test for differences between PD and noPD groups for the following variables: age, height, weight, balance ability (miniBESTest), balance confidence (ABC Scale), overall cognition (MoCA), cognitive set shifting (Trail Making Test B-A, s), years of education, N1 peak amplitudes, N1 peak latencies, and N1 peak widths. PROC TTEST in SAS was used for t-tests, including the Satterthwaite correction in cases of unequal variances. Fisher’s exact test of independence was used to test for sex differences between groups using the two-sided table probability in PROC FREQ in SAS.

#### Within-group associations

Simple linear regressions were used to test for correlations between pairs of study variables (listed below) within the PD and noPD groups separately. Parameter estimates for the regression slopes were compared against the hypothesized value 0 with two-tailed t-tests using PROC GLM in SAS. Variables that violated the assumption of normality (Shapiro-Wilk test p-values<0.05) were transformed to a normal distribution using boxcox.m in MATLAB prior to regression. Figures display original, untransformed data points with p-values and R^2^ values from the adjusted variables when appropriate. All R^2^ values are adjusted R^2^ values. Tables include Cohen’s F^2^ measure of effect size (Cohen 1992) for all simple linear regressions.

Within the noPD group, linear regressions were used to test for correlations between cortical response variables (N1 peak amplitude, latency, and width) and age, balance ability, balance confidence, TUG single task time, TUG dual task time, TUG dual task interference, overall cognition, and cognitive set shifting ability. Fisher’s exact test of independence was used to test for associations between dichotomized (median split) cortical response variables and years of education (split between N=9 with 16 or fewer years and N=10 with 18 years).

Within the PD group, linear regressions were used to test for correlations for all of the variables listed above, as well as PD duration, MDS-UPDRS-III motor symptom severity, and postural instability/gait difficulty scores. Fisher’s exact test of independence was used to test for associations between dichotomized cortical response variables and Hoehn & Yahr stage (split between N=10 at stage 2 and N=6 at stages more severe than 2), and between cortical response variables and years of education (split between N=8 with 16 or fewer years and N=8 with 18 years). Additionally, because postural instability/gait difficulty scores were distributed approximately as a negative binomial distribution, tests of association between cortical responses and postural instability/gait difficulty scores were repeated with a negative binomial regression using PROC GENMOD on the untransformed score in SAS (McKay et al. 2021).

#### Probabilistic principal components analysis

Because cortical responses were correlated with multiple measures of balance and cognitive function in the PD group (Figure 4), and because many of these variables were correlated with one another (Supplemental Information), we performed a probabilistic principal components analysis (probabilistic PCA, using ppca.m in MATLAB) to reduce the dimensionality of the covariate space to generate variables that could be entered simultaneously into a multiple regression analysis. Specifically, collinearity between variables reduces the interpretability of models that include them as separate predictors, which PCA addressed by representing variance shared across groups of variables in components that are not correlated with one another (i.e., R^2^<0.0001 for association between component loadings across participants). Probabilistic PCA is an established extension of PCA that is able to accommodate small numbers of missing values (i.e., two missing values for TUG-ST and four missing values for both TUG-DT and DTI from individuals unable to complete the tasks, as described above). The following variables were centered around zero and scaled to unit variance and entered into the probabilistic PCA: age, MDS-UPDRS-III motor symptom severity, Hoehn & Yahr stage, postural instability/gait difficulty subscores, balance ability, balance confidence, TUG single task time, TUG dual task time, TUG dual task interference, years of education, overall cognition, and cognitive set shifting ability. The first two principal components accounted for 44% (PC1) and 19% (PC2) of the total variance of the regression variables (Figure 5) and were entered into multiple regression analysis.

**Figure 3.**
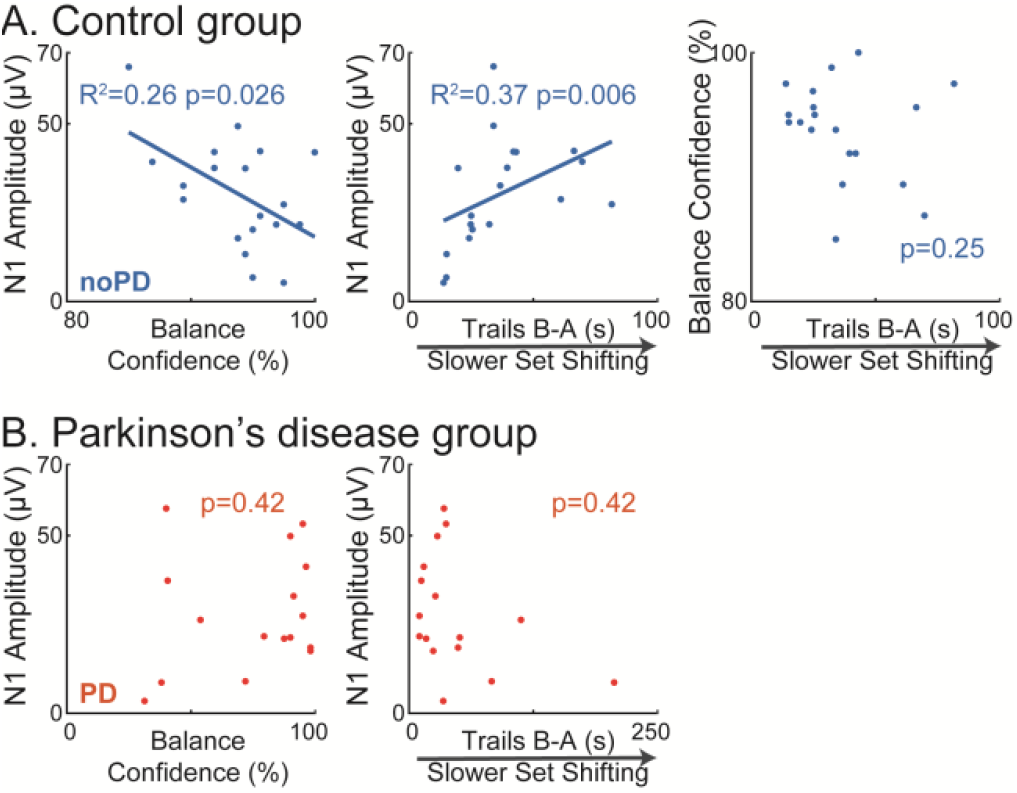
Relationships between cortical responses and clinical variables. (A) In the control group (noPD), larger N1 amplitudes were correlated with lower balance confidence and slower cognitive set shifting. Balance confidence and cognitive set shifting were not correlated with one another. (B) The group with Parkinson’s disease (PD) did not share these associations between N1 amplitude and balance confidence or cognitive set shifting. Plots show original data with statistics obtained from transformed variables when appropriate.

**Figure 4.**
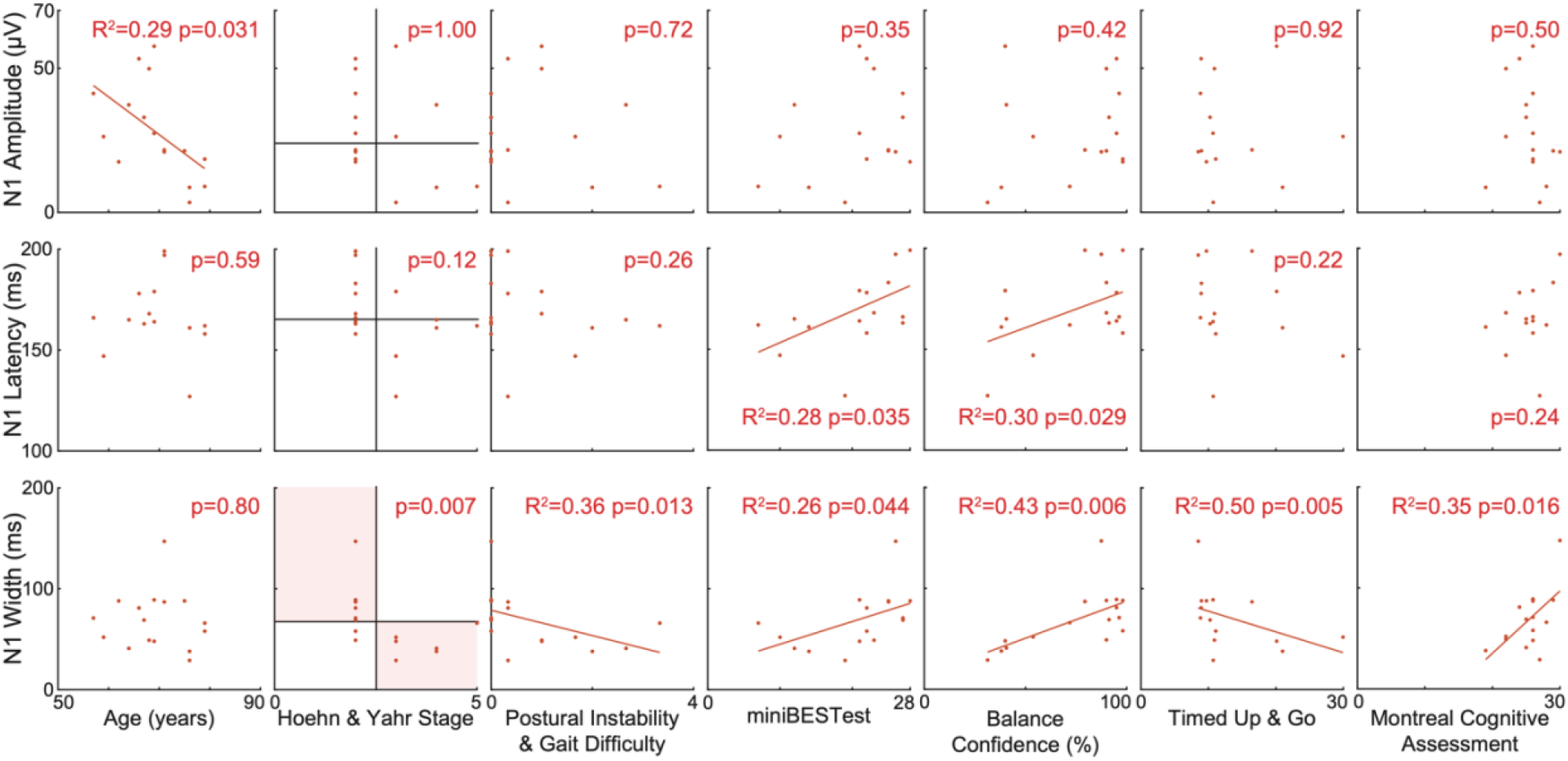
Associations between N1 measures and other variables in the PD group. Plots show original data with statistics obtained from transformed variables when appropriate.

**Figure 5.**
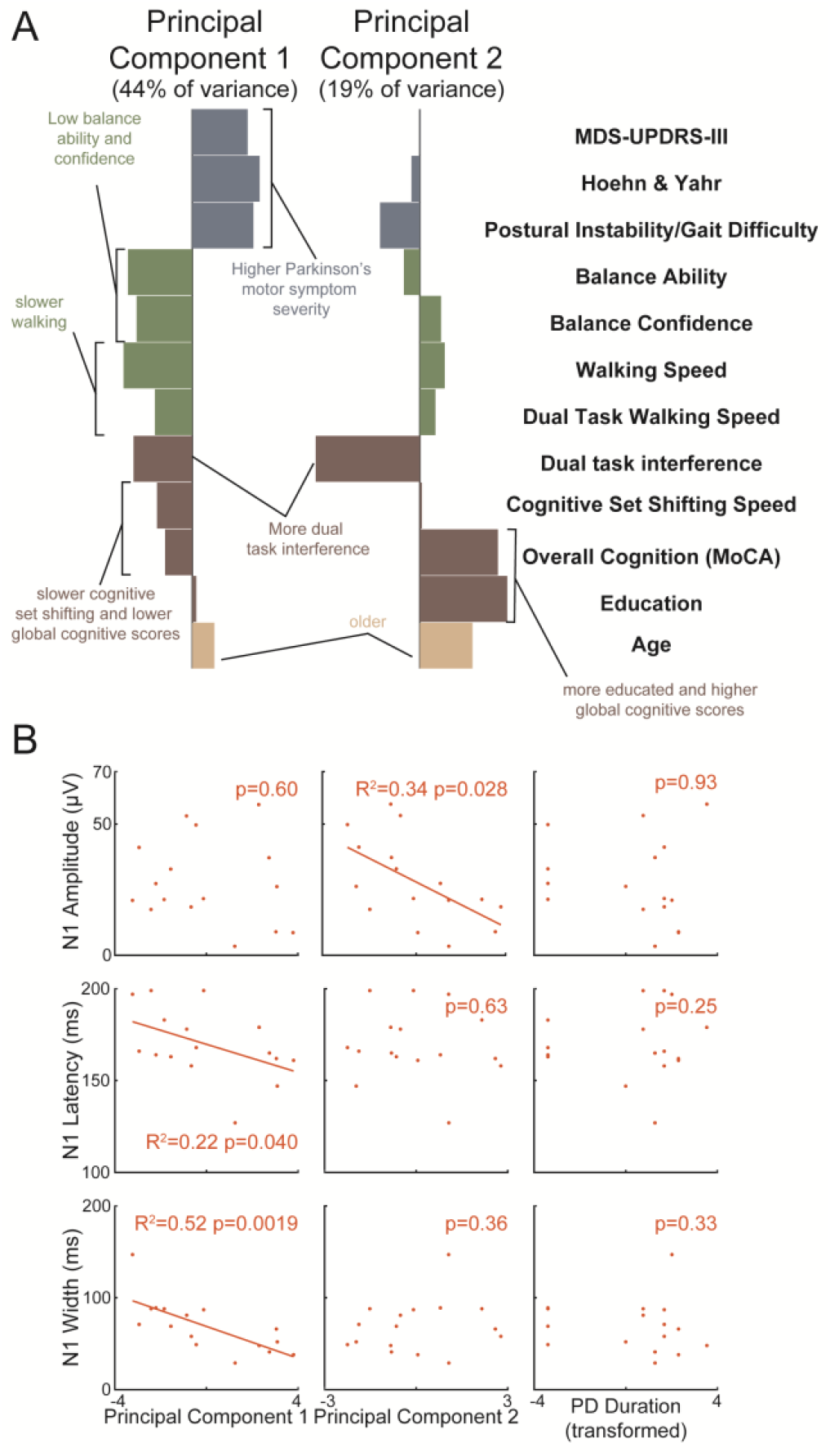
Principal components and their correlations to the N1 response in Parkinson’s disease. (A) The first two principal components (PCs) accounted for 44% and 19% of total variance, respectively. Note that PC1 has relatively more variables related to Parkinson’s disease symptom severity and balance, PC2 is dominated by dual task interference, global cognitive scores, and the age and education demographic variables on which normative data for global cognitive scores is stratified (Rossetti et al. 2011). Also note that cognitive-motor dual task interference is represented in both components. (B) Simple linear regressions are displayed along with statistics derived from the multiple regression models in which the PCs and Parkinson’s disease duration were entered as simultaneous predictors of each of the N1 measures. No outcomes differed between these simple and multiple regression models. Note that variables measured in units of time (i.e., Timed Up and Go and cognitive set shifting) have been flipped and relabeled to reflect speed (i.e., walking speed and cognitive set shifting speed) for a more intuitive representation. MDS-UPDRS: Movement Disorder Society’s Unified Parkinson’s Disease Rating scale (part III, motor symptom severity); TUG: Timed Up and Go; MoCA: Montreal Cognitive Assessment

#### Multiple linear regression analysis

Each cortical response variable (N1 peak amplitude, latency, and width) was entered into a separate multiple regression model including the two PCs and PD duration as simultaneous predictors using PROC GLM in SAS. PD duration was otherwise excluded from the principal component analysis so it could be used as a measure of PD status independent of the cognitive or motor presentation of the disease. Figures display simple linear regressions between cortical response variables and the PCs with p-values from the multiple regression analysis. No outcomes differed between these simple and multiple regression analyses. The corresponding table displays Cohen’s F^2^ value for the association between each cortical response variable and each predictor using a modified formula that considers the R^2^ value from the full model relative to the model that leaves out the variable of interest (Selya et al. 2012).

## RESULTS

### The group with Parkinson’s disease had lower balance ability and balance confidence

Participant groups (Table 1) did not differ in age (p=0.44), gender distribution (p=0.72), height (p=0.30), or weight (p=0.39). The PD group had lower balance ability (p=0.027, Cohen’s d=0.81) and balance confidence (p=0.008, d=0.98) than the noPD control group, but did not differ in overall cognition (Montreal Cognitive Assessment, p=0.41), cognitive set shifting (Trail Making Test, B-A, p=0.46), or years of education (p=0.84).

**Table 1.** Group characteristics.

|  | noPD (N=19) | PD (N=16) |
| --- | --- | --- |
| Age (years) | 71 ± 6 | 69 ± 7 |
| Gender (male/female, % female) | 13 / 6, 32% | 12 / 4, 25% |
| Height (cm) | 175 ± 10 | 171 ± 11 |
| Weight (kg) | 79 ± 16 | 85 ± 25 |
| miniBESTest ( /28) | **25 ± 2** | **21 ± 6** |
| Balance Confidence | **94 ± 4** | **75 ± 25** |
| Montreal Cognitive Assessment | 26 ± 3 | 25 ± 3 |
| Trail Making Test (B-A, s) | 61 ± 25 | 78 ± 57 |
| Education (years) | 17 ± 2 | 17 ± 2 |
| MDS-UPDRS-III |  | 31 ± 15 |
| PD Duration |  | 6 ± 3 |

Note: Bold text indicates significant group differences at $p < 0.05$ .

### Cortical N1 responses were similar between groups

Cortical N1 responses were similar between groups (Figure 2). There was a nonsignificant trend for earlier N1 peak latencies in the PD group (PD: 170±19 ms, noPD: 182±18 ms, p=0.062, d=0.63). N1 peak amplitudes (PD: 28±16 μV, noPD: 30±15 μV, p=0.66, d=0.15) and widths (full-width half-maximum, PD: 69±29 ms, noPD: 85±39 ms, p=0.17, d=0.47) were similar between groups.

### In the control group, N1 amplitudes were associated with higher balance confidence and lower cognitive set shifting ability

In the noPD group, larger N1 amplitudes were correlated with lower balance confidence (Figure 3A, p=0.026, R^2^=0.26, F^2^=0.35). As reported previously (Payne et al. 2021), larger N1 amplitudes were correlated with lower cognitive set shifting ability (p=0.006, R^2^=0.37, F^2^=0.57). Balance confidence was not associated cognitive set shifting ability (p=0.25). There was a trend for an association between more years of education and wider N1 peaks (p=0.070). N1 amplitude, latency, and width were not associated with any other tested variables in the noPD group (Table 2). These associations were not observed in the PD group (Table 3 and shown in Figure 3B for comparison).

**Table 2.** Associations between cortical responses and other variables in the control group.

| noPD group | N1 Amplitude |  | N1 Latency |  | N1 Width |  |
| --- | --- | --- | --- | --- | --- | --- |
|  | F<sup>2</sup> | p | F<sup>2</sup> | p | F<sup>2</sup> | p |
| Age | 0.05 | 0.378 | 0.07 | 0.284 | 0.05 | 0.388 |
| miniBESTest | 0.00 | 0.928 | 0.02 | 0.551 | 0.05 | 0.379 |
| Balance Confidence | **0.35** | **0.026** | 0.01 | 0.753 | 0.09 | 0.226 |
| TUG-Single Task | 0.19 | 0.091 | 0.05 | 0.367 | 0.09 | 0.238 |
| TUG-Dual Task | 0.10 | 0.215 | 0.02 | 0.605 | 0.00 | 0.959 |
| Dual Task Interference | 0.04 | 0.448 | 0.04 | 0.419 | 0.01 | 0.676 |
| Education | - | 0.180 | - | 0.370 | - | 0.070 |
| Montreal Cognitive Assessment | 0.02 | 0.580 | 0.02 | 0.555 | 0.25 | 0.057 |
| Cognitive Set Shifting | **0.57** | **0.006** | 0.12 | 0.175 | 0.03 | 0.500 |

Note: Bold text indicates significant associations at $p < 0.05$ . Cohen's $F^2 > 0.35$ indicates a large effect and $F^2 > 0.15$ indicates a medium effect. TUG: Timed Up and Go

**Table 3.**
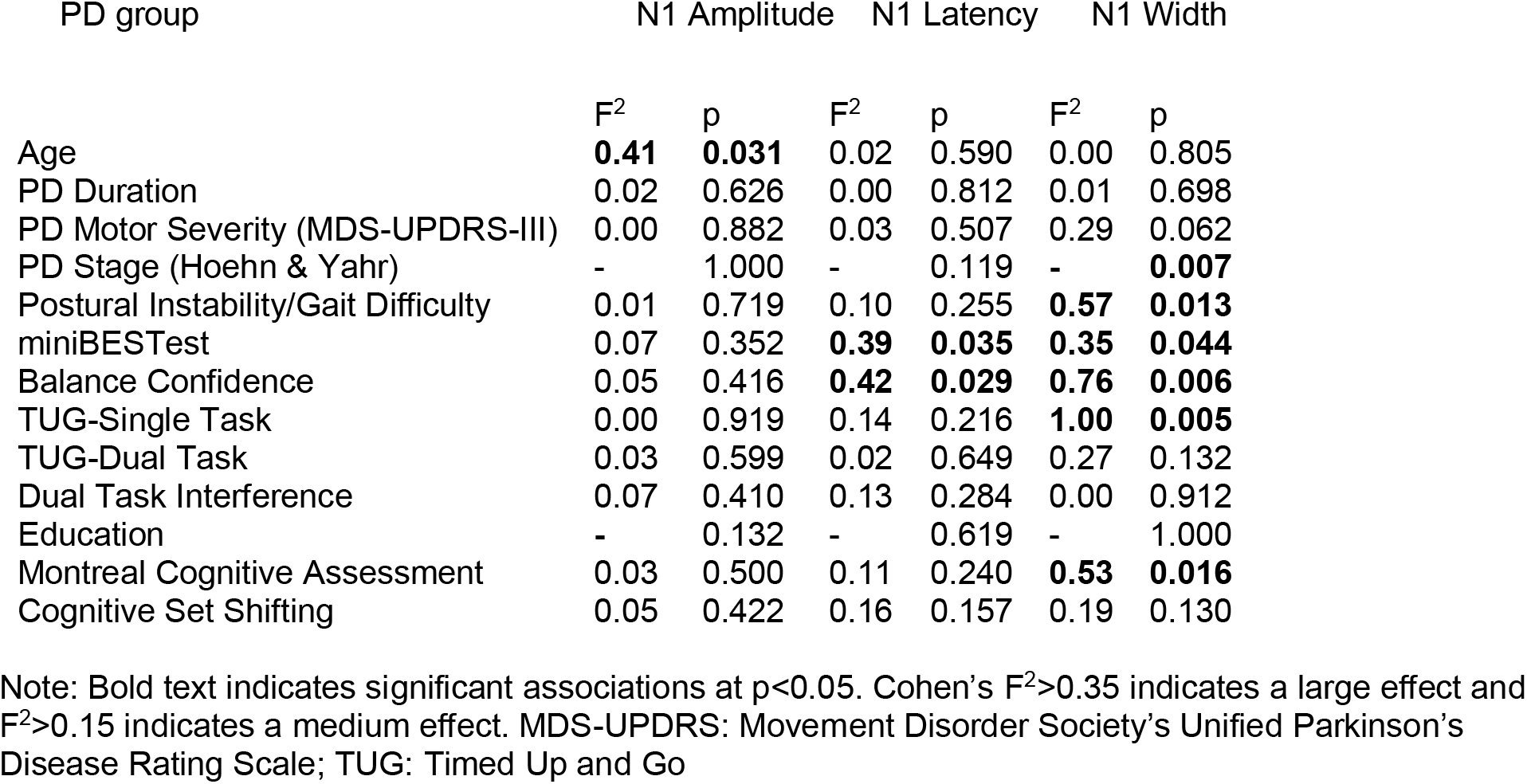
Associations between cortical responses and other variables in the group with Parkinson’s disease.

### N1s were associated with multiple overlapping measures of balance and cognitive function in the group with Parkinson’s disease

In the PD group, cortical N1 responses were associated with overlapping measures of balance and cognitive function (Table 3, Figure 4). Larger N1 amplitudes were correlated with younger age (p=0.031, R^2^=0.29, F^2^=0.41). Longer N1 latencies were correlated with higher clinical balance ability (p=0.035, R^2^=0.28, F^2^=0.39) and higher balance confidence (p=0.029, R^2^=0.30, F^2^=0.42). Narrower N1 peak widths were associated with more severe Hoehn & Yahr disease stages (Fisher’s exact test, p=0.007), more severe postural instability/gait difficulty subscores (linear regression p=0.013, R^2^=0.36, F^2^=0.57, negative binomial regression p=0.033), lower mobility (slower single task TUG, p=0.005, R^2^=0.50, F^2^=1.00), lower balance ability (p=0.044, R^2^=0.26, F^2^=0.35), lower balance confidence (p=0.006, R^2^=0.43, F^2^=0.76), and lower overall cognitive ability (Montreal Cognitive Assessment, p=0.016, R^2^=0.35, F^2^=0.53). The cortical N1 responses were not associated with the other tested variables in the PD group (Table 3).

Principal components analysis was applied to the dataset to reduce the number of comparisons and to account for covariation between the tested variables. Correlations between all pairs of tested variables are reported in Supplemental Information. The first two principal components accounted for 44% and 19% of the variance of the dataset (Figure 5).

The N1 amplitudes were associated with PC2 (Table 4, Figure 5), while the N1 peak latency and peak width were associated with PC1. In the multiple regression model, larger N1 amplitudes were correlated with lower component loadings on PC2 (representing lower dual-task interference on walking, lower scores on a global measure of cognitive function, younger age, and fewer years of education, p=0.028, F^2^=0.52), but not PC1 (p=0.60) or Parkinson’s disease duration (p=0.93) included in the same model. Shorter N1 peak latencies were correlated with PC1 (representing higher PD severity, lower balance function and confidence, slower walking speed, greater dual-task interference on walking, and slower cognitive set shifting, p=0.040, F^2^=0.44) but not PC2 (p=0.63) or Parkinson’s disease duration (p=0.25). Narrower N1 peak widths were also correlated with PC1 (p=0.002, F^2^=1.31) but not PC2 (p=0.36) or Parkinson’s disease duration (p=0.33).

**Table 4.** Associations between cortical responses and principal components in the group with Parkinson’s disease.

|  | N1 Amplitude |  | N1 Latency |  | N1 Width |  |
| --- | --- | --- | --- | --- | --- | --- |
| | $F^2$ | p | $F^2$ | p | $F^2$ | p |
| PC1 | 0.02 | 0.601 | **0.44** | **0.040** | **1.31** | **0.002** |
| PC2 | **0.52** | **0.028** | 0.02 | 0.628 | 0.07 | 0.363 |
| PD Duration | 0.00 | 0.929 | 0.12 | 0.245 | 0.08 | 0.334 |

Note: Statistics refer to multiple regression models where the three row variables are entered as simultaneous predictors of the corresponding column variable. Bold text indicates significant associations at $p<0.05$ . Cohen's $F^2>0.35$ indicates a large effect and $F^2>0.15$ indicates a medium effect.

## DISCUSSION

This is the first paper to compare the balance perturbation-evoked cortical N1 response between people with and without Parkinson’s disease. N1 responses were similar in amplitude, latency, and peak width between groups, but were associated with different balance- and cognitive-related measures in older adults with versus without Parkinson’s disease. We previously reported that larger N1 responses were associated with lower cognitive set shifting ability in older adults (Payne et al. 2021), and we now add with the present study that the larger N1 responses are also associated with lower balance confidence in the same cohort. However, N1 responses in the group with Parkinson’s disease did not share these simple correlations with cognitive set shifting or balance confidence, but rather were associated with multiple overlapping measures related to balance and cognitive function. Within the Parkinson’s disease group, distinct features of the N1 responses were associated with two principal components representing variance shared across groups of variables. Larger N1 amplitudes in the group with Parkinson’s disease were correlated with a principal component representing lower scores on a global measure of cognition, lower dual-task interference, younger age, and fewer years of education. Earlier and narrower N1 peaks were correlated with a principal component representing greater parkinsonian motor symptom severity, greater balance impairment, lower balance confidence, slower walking speed, and greater dual-task interference. Our results show that individual differences in balance and cognitive function in Parkinson’s disease are reflected in the N1 response, which may reflect individual differences in activation of the various cortical areas that synchronize during the N1 response. A better understanding of the neural mechanisms related to balance and cognitive impairments could facilitate the development of more targeted rehabilitation for individuals with co-occurring balance and cognitive impairments.

The lack of differences in the cortical N1 response at the group level suggests there is no specific effect of Parkinson’s disease or dopamine depletion on the N1 response. The similarity of the perturbation-evoked N1 response between groups is a clear contrast from the dopamine-dependent error-related negativity, which is robustly reduced in amplitude even at early stages of Parkinson’s disease, including shortly after diagnosis, before initiation of dopaminergic treatment (reviewed in Seer et al. 2016). Whereas the error-related negativity is time-locked to the behavioral response, stimulus-locked potentials in cognitive tasks such as the P3, mismatch negativity, and conflict-N1 are not consistently altered in Parkinson’s disease, except possibly within select subtypes of cognitive impairment (Seer et al. 2016), which the present study did not distinguish between. However, movement-related cortical potentials that precede voluntary movement (Cunnington et al. 1995; Oishi et al. 1995; Shibasaki et al. 1978) and the initial negativity in somatosensory-evoked potentials after median nerve stimulation (Bostantjopoulou et al. 2000; Rossini et al. 1989) are reduced in Parkinson’s disease, so it is unclear why the perturbation-evoked N1 seems to be selectively preserved. A slow negative deflection called the contingent negative variation, resembling the initial component of movement-related cortical potentials, precedes the N1 response when balance perturbations are predictable (Mochizuki et al. 2010; Mochizuki et al. 2009b), and this also appears to be preserved in amplitude in Parkinson’s disease, although lacking associated preparation-related changes in behavior (Smith et al. 2012). Although we would expect an attenuated N1 in Parkinson’s disease based on the most comparable cognitive, motor, and sensory potentials, the extension of individual differences within healthy populations would suggest an enhancement of the N1 in Parkinson’s disease. Specifically, N1 amplitudes are larger in those with lower balance ability among young adults (Payne and Ting 2020a), and in those with lower balance confidence among older adults, but the group differences of lower balance ability and lower balance confidence in Parkinson’s disease were not accompanied by group differences in N1 amplitudes. While we cannot rule out the possibility that an enhanced N1 due to lower balance ability and balance confidence is counteracted by an attenuation of the N1 response due to dopamine depletion in Parkinson’s disease, the present data provide no evidence to suggest that the cortical N1 response depends on dopamine function, or on the basal ganglia and brainstem centers that are affected by Parkinson’s disease.

Larger N1 amplitudes were associated with two factors known to predict falls in older adults. Prior work largely focused on within-subjects effects, which can demonstrate causality, but do not guarantee the same information in terms of individual differences (Hajcak et al. 2017), which must be established before any potential biomarker can inform clinical decision-making. The present finding of larger N1 amplitudes in older adults with lower balance confidence seems to extend work showing N1 amplitudes increase in more threatening contexts (Adkin et al. 2008; Mochizuki et al. 2010). That is, larger N1 amplitudes may indicate greater concern about falling both within and between individuals. Although balance confidence predicts falls in older adults (Cleary and Skornyakov 2017), predictive validity of the N1 has not been investigated in any context. Similarly, as previously reported (Payne et al. 2021), N1 amplitudes are larger in individuals with slower cognitive set shifting, which is a measure of executive function that also predicts falls in older adults (Muir et al. 2012). Although not a direct parallel, this finding adds another line of evidence connecting the N1 to cognitive processing, along with within-subjects effects of surprise (Adkin et al. 2008; Mochizuki et al. 2010) and distraction (Little and Woollacott 2015; Quant et al. 2004b). That is, cortical processes involved in cognitive set shifting may overlap with those influencing the N1 response when distracted or surprised, or there may be multiple mechanisms though which changes in cognitive processing influence the N1. It would therefore be interesting to test whether the magnitude of the dual-task effect on the N1 response is associated with individual differences in cognitive ability. Although prior findings of larger N1 amplitudes in young adults with worse balance (Payne and Ting 2020a) did not extend to the present cohort of older adults, the clinical balance scores displayed a ceiling effect with scores clustered near the top of the range in our unimpaired older adult population (Payne et al. 2021). While the N1 could relate to a more challenging metric of balance ability in older adults, this could also reflect a difference in the N1 response across the lifespan. This also suggests the N1 response differs from other measures of brain activity during balance recovery, such as beta frequency (13-30 Hz) power, which is associated with balance ability in young adults (Ghosn et al. 2020) and clinical balance ability in older adults (Palmer et al. 2021). In any case, it would be interesting to investigate longitudinal predictive validity of the N1 response, much like work demonstrating prefrontal oxygenation during dual task walking predicts falls (Verghese et al. 2017) and resting state low frequency power predicts cognitive decline (Caviness et al. 2015; Cozac et al. 2016).

N1 responses were associated with multiple overlapping measures related to balance and cognition in the group with Parkinson’s disease, further suggesting the N1 response may reflect mechanisms related to both balance and cognitive impairments. Through our second principal component, larger N1 amplitudes appear to be associated with lower scores on a global measure of cognition, fewer years of education, less dual task interference on walking, and younger age. However, this should be interpreted with caution as only the association with age was supported by the simple correlations, which also suggested that lower global cognitive scores were instead related to narrower N1 widths. In contrast, associations between temporal features of the N1 response and the first principal component, representing Parkinson’s disease motor symptom severity, lower balance ability and confidence, and lower mobility were largely supported by the simple correlations. Specifically, lower balance ability and confidence were correlated with both earlier and narrower N1s, while narrower N1s were additionally correlated with slower walking speed and more advanced Parkinson’s disease balance disability. Although there were no simple correlations between the N1 response and cognitive set shifting in the group with Parkinson’s disease, the inclusion of cognitive set shifting in the principal component that represented most of the balance-related measures is consistent with prior work linking cognitive set shifting ability to falls in older adults with and without Parkinson’s disease (McKay et al. 2018; Muir et al. 2012). Additionally, the other principal component represented an association between lower postural instability/gait difficulty scores and higher global cognitive function, consistent with longitudinal work showing that postural instability/gait difficulty develops in tandem with accelerated cognitive decline in Parkinson’s disease (Alves et al. 2006). The association between temporal features of the N1 response and balance ability in the group with Parkinson’s disease is in contrast to the association between N1 amplitude and balance ability in young adults (Payne and Ting 2020a), but this is not the first study to link motor ability to temporal features of the N1 response (Duckrow et al. 1999). Despite different relationships across populations, the present results suggest that the N1 response reflects neural processes related to both balance and cognitive function, and thus a better understanding of the underlying neural mechanisms could provide insight into the associations between balance and cognitive decline in Parkinson’s disease.

We speculate that the N1 response reflects neural processes at the intersection of balance and cognitive function that could explain relationships between balance and cognitive impairments and their overlapping responses to treatment in aging populations. Although we are unable to separate and localize the underlying neural sources due to our limited electrode set, studies in young adults suggest multiple neural sources synchronize during the N1 response (Peterson and Ferris 2018; 2019; Varghese et al. 2019). It is possible that different relationships between the N1 and the various individual difference measures across populations reflect differences in the relative contributions of the multiple neural sources underlying the N1 response across populations. Additionally, the appearance of multiple component peaks in older populations (Dimitrov and Gatev 2001; Duckrow et al. 1999; Payne et al. 2021) could arise through loss of coordination between these underlying neural sources, manifested as reduced synchrony, coherence, or phase alignment of theta oscillatory power across the underlying sources, or as suggested by Duckrow et al. 1999, reduced myelination with aging could cause some discrete subcomponent to be delayed relative to others. It is possible that changes in the interactions between neural processes involved in balance and cognitive function could underlie associations between balance and cognitive decline in aging populations (Allcock et al. 2009; Camicioli and Majumdar 2010; Gleason et al. 2009; Herman et al. 2010; Mak et al. 2014; Mirelman et al. 2012), and might explain reciprocal crossover benefits between balance and cognitive rehabilitation (Hagovska and Olekszyova 2016; Kraft 2012; Manor et al. 2018; Smith-Ray et al. 2015). If the N1 response reflects neural processes at the intersection of balance and cognitive function, a deeper understanding of the underlying mechanisms could facilitate the development of more targeted rehabilitation for individuals with co-occurring balance and cognitive impairments.

## Acknowledgements

Study data were collected and managed using a Research Electronic Data Capture (REDCap) database hosted at Emory University (Harris et al. 2019; Harris et al. 2009).

